# Multimorbidity and healthcare utilization: A register-based study in Denmark

**DOI:** 10.1101/574293

**Authors:** Anne Frølich, Nermin Ghith, Michaela Schiøtz, Ramune Jacobsen, Anders Stockmarr

## Abstract

**Background:** People with multimorbidity have reduced functional capacity, lower quality of life, and higher mortality rates and use healthcare resources more intensively than healthy people or those with a single chronic condition. The aim of this study was to explore associations between multimorbidity and use of healthcare services and the impact of socioeconomic status on utilization of hospitalizations and bed days.

**Methods:** The study population included all individuals aged 16 years and older who lived in the Capital Region of Denmark on January 1st, 2012. Data on chronic conditions, use of healthcare services and demographics were obtained from Danish national administrative and health registries. Zero-inflated models were used to calculate anticipated annual use of hospitalizations and bed days.

**Findings:** The study population comprised 1,397,173 individuals; the prevalence of multimorbidity was 22%. Prevalence was inversely related to educational attainment. For people with multimorbidity, utilization of hospitalizations and bed days increased approximately linearly with the number of chronic conditions. However, a steep increase in utilization of bed days was observed between five and six or more chronic conditions. An educational gradient in hospitalization rates and use of bed days was observed regardless of the number of chronic conditions. Educational attainment was strongly associated with healthcare utilization.

**Conclusion:** Multimorbidity was associated with a significant increase in utilization of all healthcare services in Denmark. In addition, a socioeconomic gradient was observed in utilization of hospitalizations and bed days.

## Introduction

Multimorbidity is often defined as the coexistence of two or more chronic conditions in the same person [1, 2]. People with multimorbidity have decreased functional competence [3], lower quality of life [4], higher mortality rates [5] and use healthcare resources more intensively than healthy people or those with just one chronic condition [6]. Most diseases and consequences of poor health are unequally distributed among socioeconomic population groups, and socioeconomic differences are obvious in the prevalence and consequences of multimorbidity [7, 8]. The prevalence of multimorbidity is increasing internationally. The overall prevalence of multimorbidity in Ontario, Canada increased in nearly all age groups, reflecting a 40% total increase between 2003 and 2009 from 17.4% to 24.3% [9]. A study from the Netherlands reported an increase in multimorbidity prevalence from 12.7% to 16.2% between 2004 and 2011 [10]. An American study showed that the prevalence of multimorbidity rose between 2000 and 2010 from 22% to 30%, a trend that was most pronounced among people younger than age 65 [11]. Expected continuing increases in the prevalence of multimorbidity are recognized as a major public health and healthcare challenge for modern societies [9].

To understand the healthcare challenges associated with multimorbidity, the impact of multimorbidity on healthcare utilization must be carefully assessed [12, 13]. However, detailed knowledge about how multimorbidity affects healthcare utilization is incomplete. A systematic review identified 35 studies investigating relationships between multimorbidity and healthcare utilization, healthcare costs, or both. All included studies showed a positive correlation between multimorbidity and at least one aspect of utilization (physician visits and hospital care) and costs (medications, out-of-pocket spending, and total healthcare costs) for elderly populations [14]. The included studies were from the United States (23), Europe (5), Canada (4), Asia (2), and Australia (1). Many of these study populations were large enough to enable sophisticated statistical analyses, and four were large cross-sectional studies that included 1.13 million to 1.65 million people [15-17]. Regrettably, these large studies had varying inclusion criteria, and none reported sociodemographic data that are necessary for exploring possible associations between multimorbidity, healthcare utilization, and sociodemographic status. Synthesis of the studies was not possible due to ambiguous definitions and measurements of multimorbidity, as well as a multitude of healthcare utilization outcomes.

After publication of this systematic review, several large-scale studies [18-22] exploring the relationship between healthcare utilization and the number of chronic conditions demonstrated that additional factors affecting utilization include age [18, 19], gender [18, 19], impaired activities of daily life [20], and socioeconomic status [19]. Furthermore, the impact of multimorbidity on healthcare utilization has been shown to differ across individual factors, disease combinations, healthcare systems, and regions [22, 23].

The Danish healthcare system is based on universal coverage and principles of free and equal access to healthcare for all citizens. Information on the impact of multimorbidity on healthcare utilization in the Danish healthcare system is sparse. To the best of our knowledge, only a single study to date has demonstrated that hospitalizations, use of bed days and general practitioner (GP) visits, were significantly higher for patients with multimorbidity, compared with those who had no chronic conditions [24].

The aim of this study was to explore associations between multimorbidity and healthcare utilization and the impact of socioeconomic status on utilization of hospitalizations and bed days. The structure and content of Danish healthcare registers provide a unique opportunity to quantify how these variables interact [25, 26], which we explored in a large-scale, cross-sectional, regional, register-based population study.

## Methods

### Study population and data sources

The study population included all individuals aged 16 years and older who lived in the Capital Region of Denmark on January 1st, 2012. The included 1,397,173 individuals represented approximately one-third of the entire Danish adult population. Data on chronic conditions, use of healthcare services, and demographics, including gender, age, and educational attainment, were obtained from Danish national administrative and health registries: the Danish National Patient Registry [27], the Danish National Prescription Registry [28], the Danish National Health Service Registry [29], and The National Diabetes Registry [30]. All data obtained from registries were merged at the level of individuals using their unique social security numbers. However, national registries do not provide data on conditions diagnosed in the primary care sector. In addition to data on diagnoses of chronic conditions available from the secondary healthcare sector, diagnostic algorithms developed by the Research Center for Prevention and Health at Glostrup University Hospital were used to identify primary sector diagnoses of 16 chronic conditions of interest for the entire population (Table 1) [6]. Details about the diagnostic algorithms are provided elsewhere [7]. Multimorbidity was defined as two or more chronic conditions occurring simultaneously in the same person.

**Table 1.**
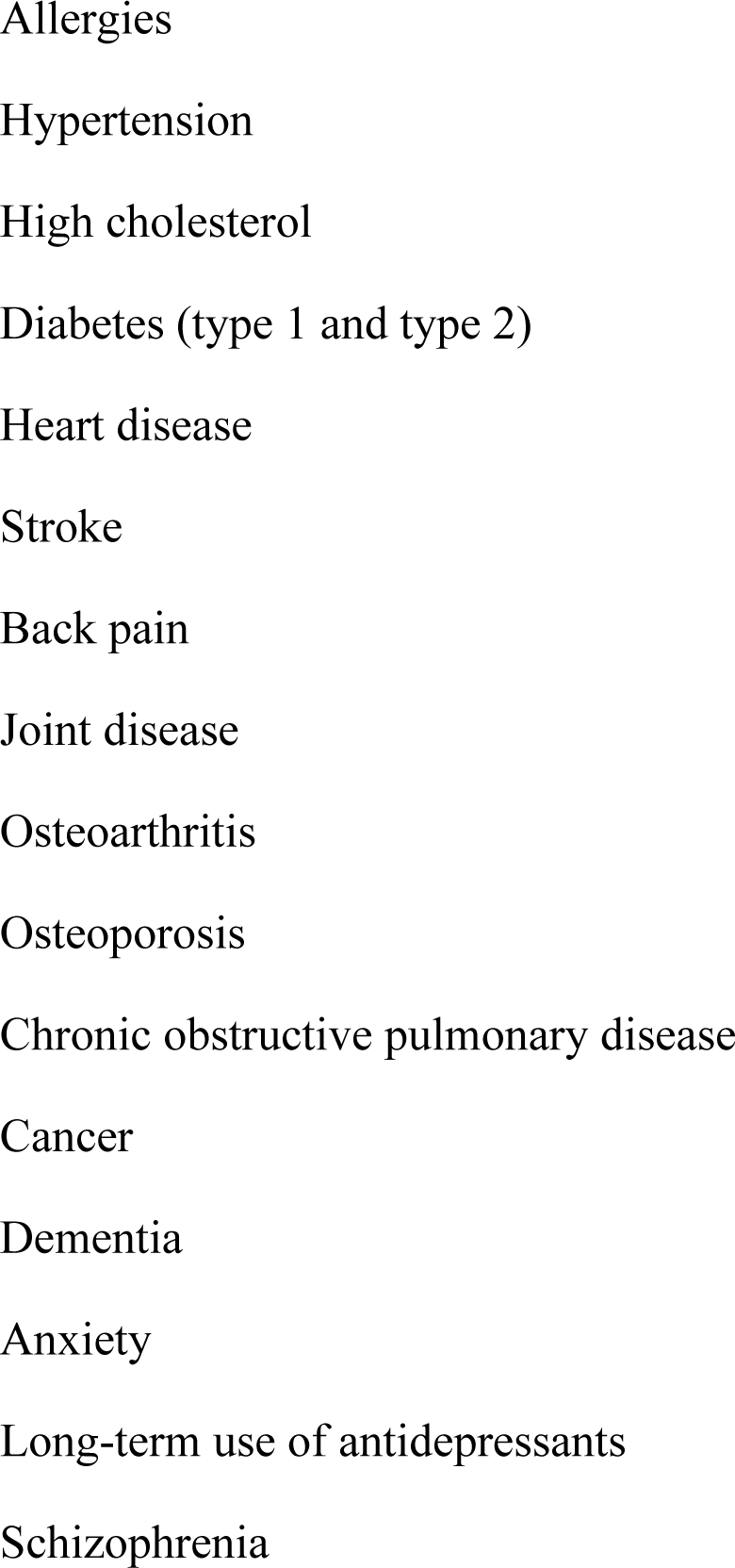
Chronic conditions included in definition of multimorbidity (N = 16)

Because data on direct costs could not be linked to the study data, we identified utilization of hospitalizations and bed days, which are among the most expensive services in healthcare, as proxies for direct costs. Socioeconomic status was defined as highest educational achievement and grouped into four categories according to the length of education: none (primary school) (≤10 years), short (11-14 years), medium (14-17 years), and long (≥17 years) [31]. Finally, we also recorded healthcare utilization for emergency department visits, outpatient visits, GP visits, out-of-hour GP visits, yearly control visits in general practice, and specialist visits during 2012.

## Statistical analysis

Descriptive statistics were calculated for gender, age, and educational attainment by the number of chronic conditions. Means for each type of healthcare utilization were calculated. Logistic regression was used to calculate odds ratios (ORs) for healthcare utilization for individuals with multimorbidity versus individuals with none and with one chronic condition. ORs were calculated in both a raw form and adjusted for gender, age, and educational attainment.

As cost data was not linked to the present dataset we chose to investigate the effect of socioeconomic status based on the proxies for general healthcare utilization: hospitalizations and bed days. Hospitalizations and bed days were adjusted for emergency visits, out-patient visits GP visits, out-of-hours GP visits, yearly controls in general practice, private specialist visits, number of conditions, age, gender, cohabitation status, education attainment, and employment status.

This was accomplished by applying zero-inflated models to both proxies. The models were used to calculate anticipated annual use of hospitalizations and bed days within educational attainment groups, separately for each number of chronic conditions.

The decision to use zero-inflated models was prompted by the fact that many individuals do not use bed days or hospitalization [32]. To counter extreme values for some covariates, squared effects of all numerical covariates were included as explanatory variables. For the partial model (1) below, a squared effect of GP visits was not included due to convergence problems. For the partial models of the form (2) below, quadrupled effects of ambulatory visits and out-of-hours GP visits were included. The zero-inflated model was applied as follows.

First, the probability *p* of having at least one bed day/one hospitalization was modeled with a logistic regression model

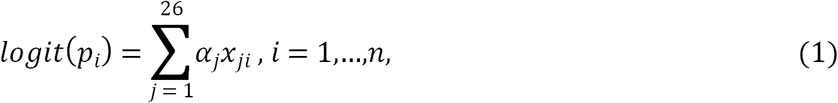

where *p*_*i*_ is the probability of one or more hospitalizations or bed days for an individual *i* and *x*_*j*_ is the *j*th covariate derived from the specified explanatory variables. Subsequently, numbers of bed days and hospitalizations *y* were similarly modeled with a log-linear model,

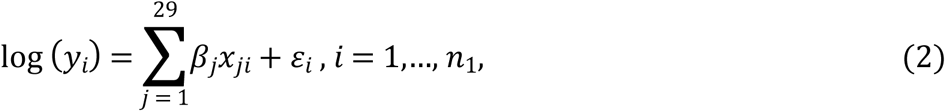

where *n*_1_ ≤ *n* is the number of individuals with at least one bed day or hospitalization. In the models (2), two observations of bed days and three observations of hospitalizations were omitted as outliers for the analysis of bed days and hospitalizations, respectively.

For scenarios in which a specific number of conditions and a specific level of education were assumed, the linear predictors 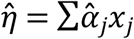 and 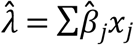 were calculated, with all other covariates kept at the empirical mean for the group with the specified number of conditions and level of education. Estimated responses were found as 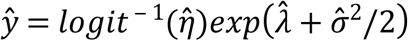, where *σ*^2^ is the variance of 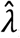, thus combining models (1) and (2). All analyses were carried out using R software version 3.5.0 (34).

## Ethics approval

Approval to conduct the study was obtained from the Danish Data Protection Agency. No informed consent was required.

## Results

Among 1,397,173 individuals included in the study population, approximately half (720,885; 52%) were women, the majority (927,568; 66%) had none or short education, and the prevalence of multimorbidity was 22% (301,757). Table 2 shows the distribution of gender, age, and educational attainment by the number of chronic conditions. The prevalence of both one and two or more chronic conditions was significantly higher for women than for men. Overall, multimorbidity was highest among individuals 65-84 years old, followed by those who were 45-64 years old. Educational attainment was inversely related to multimorbidity.

**Table 2.** Distributions of the study population (N = 1,397,173) by gender, age, and educational attainment and number of chronic conditions, N (%)

|  |  | Number of chronic conditions |  |  |
| --- | --- | --- | --- | --- |
| | | 0 | 1 | $\geq 2$ |
|  |  | N = 809,920 | N = 285,496 | N = 301,757 |
| Gender |  |  |  |  |
|  | Male | 418,105 (62) | 127,591 (19) | 130,592 (19) |
|  | Female | 391,815 (54) | 157,905 (22) | 171,165 (24) |
| Age in years |  |  |  |  |
|  | 16-24 | 170,486 (86) | 25,768 (13) | 1,770 (1) |
|  | 25-44 | 385,540 (77) | 89,216 (18) | 23,557 (5) |
|  | 45-64 | 205,947 (48) | 111,147 (26) | 110,949 (26) |
|  | 65-84 | 45,529 (19) | 53,334 (22) | 140,001 (59) |
|  | >84 | 2,418 (7) | 6,031 (18) | 25,480 (75) |
| Education |  |  |  |  |
| | None ($\leq 10$ years) | 185,347 (53) | 69,779 (20) | 93,702 (27) |
|  | Short (11-14 years) | 325,241 (56) | 123,439 (22) | 130,420 (23) |
|  | Medium (15-16 years) | 132,531 (59) | 49,576 (22) | 41,536 (19) |
| | Long ($\geq 17$ years) | 107,578 (65) | 34,035 (21) | 22,174 (14) |
|  | Missing | 59,233 (72) | 8,667 (11) | 13,925 (17) |

Mean rates of healthcare utilization among people with multimorbidity were much higher than among people with no chronic conditions, by a factor of 1.73 to 9.67, depending on the type of utilization (Table 3). When comparing people with multimorbidity and those with one chronic condition, rates of healthcare utilization were 1.44 to 4.00 times higher for those with multimorbidity. In both comparisons, the largest between-group difference was for yearly control visits in general practice. Unadjusted and adjusted ORs for all types of healthcare utilization were significantly higher (p<0.001) among people with multimorbidity, compared with people with zero or one chronic condition. Unadjusted ORs for healthcare utilization for people with multimorbidity were 1.65 to 11.76 times higher than for people with no chronic conditions and 1.31 to 3.15 times higher than for people with one chronic condition (Table 3). Adjusted ORs for healthcare utilization for people with multimorbidity were 1.86 to 6.70 times higher than for people with no chronic conditions and 1.44 to 2.94 times higher than for people with one chronic condition (Table 3).

**Table 3.**
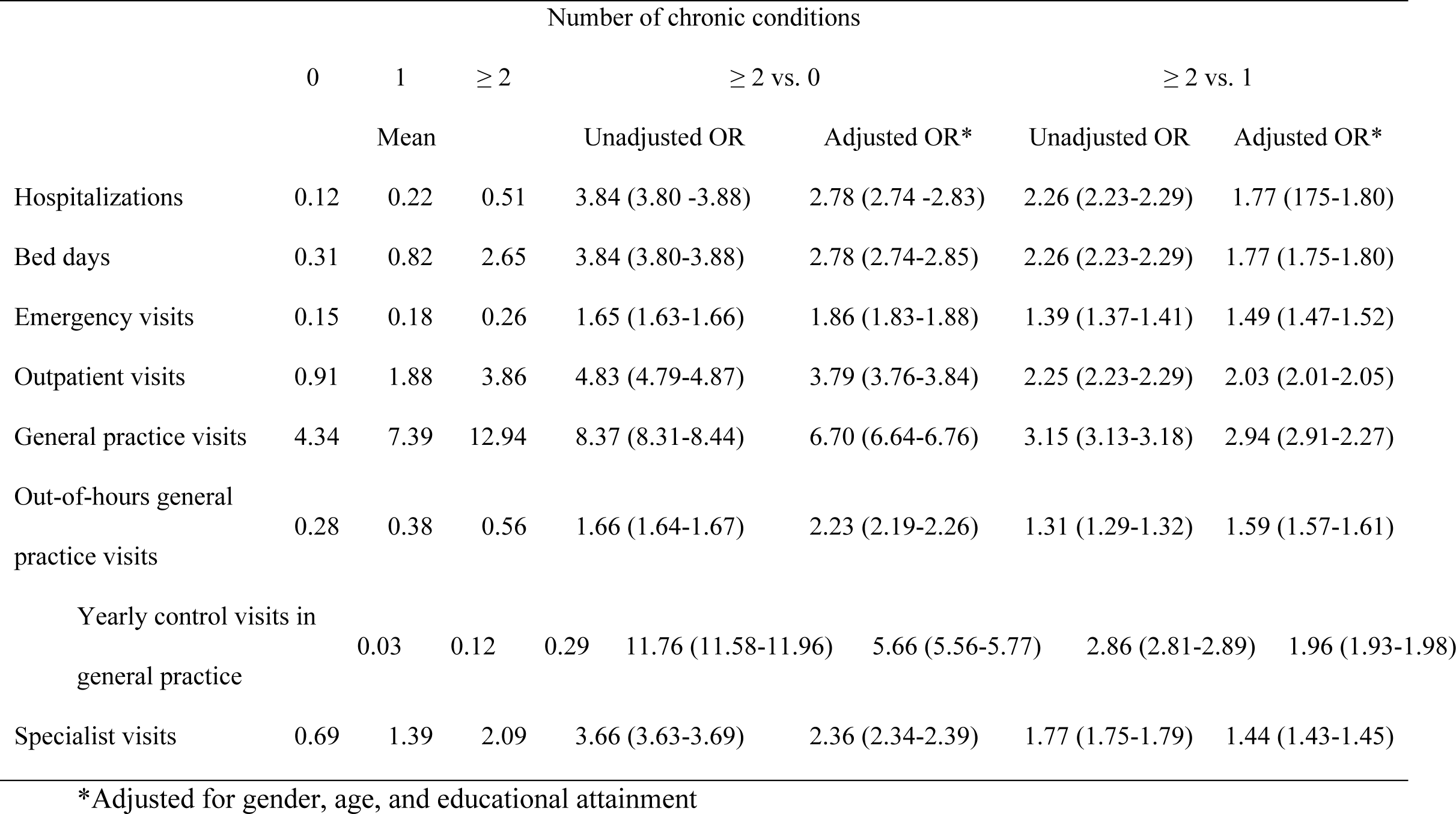
Mean and odds ratios (ORs) for healthcare utilization among individuals with 0, 1, and ≥ 2 chronic conditions

|  | Number of chronic conditions |  |  |  |  |  |  |
| --- | --- | --- | --- | --- | --- | --- | --- |
| | 0 | 1 | $\geq 2$ | $\geq 2$ vs. 0 | | $\geq 2$ vs. 1 | |
|  | Mean |  |  | Unadjusted OR | Adjusted OR* | Unadjusted OR | Adjusted OR* |
| Hospitalizations | 0.12 | 0.22 | 0.51 | 3.84 (3.80-3.88) | 2.78 (2.74-2.83) | 2.26 (2.23-2.29) | 1.77 (1.75-1.80) |
| Bed days | 0.31 | 0.82 | 2.65 | 3.84 (3.80-3.88) | 2.78 (2.74-2.85) | 2.26 (2.23-2.29) | 1.77 (1.75-1.80) |
| Emergency visits | 0.15 | 0.18 | 0.26 | 1.65 (1.63-1.66) | 1.86 (1.83-1.88) | 1.39 (1.37-1.41) | 1.49 (1.47-1.52) |
| Outpatient visits | 0.91 | 1.88 | 3.86 | 4.83 (4.79-4.87) | 3.79 (3.76-3.84) | 2.25 (2.23-2.29) | 2.03 (2.01-2.05) |
| General practice visits | 4.34 | 7.39 | 12.94 | 8.37 (8.31-8.44) | 6.70 (6.64-6.76) | 3.15 (3.13-3.18) | 2.94 (2.91-2.27) |
| Out-of-hours general practice visits | 0.28 | 0.38 | 0.56 | 1.66 (1.64-1.67) | 2.23 (2.19-2.26) | 1.31 (1.29-1.32) | 1.59 (1.57-1.61) |
| Yearly control visits in general practice | 0.03 | 0.12 | 0.29 | 11.76 (11.58-11.96) | 5.66 (5.56-5.77) | 2.86 (2.81-2.89) | 1.96 (1.93-1.98) |
| Specialist visits | 0.69 | 1.39 | 2.09 | 3.66 (3.63-3.69) | 2.36 (2.34-2.39) | 1.77 (1.75-1.79) | 1.44 (1.43-1.45) |

\*Adjusted for gender, age, and educational attainment

Among people with multimorbidity, utilization of hospitalizations increased approximately linearly with the number of chronic conditions. In Fig. 1A, hospitalization rates are indicated with a black line; the reference regression line in red has a slope equal to the mean number of hospitalizations across the number of chronic conditions. The similarity between observed rates and the regression line indicates that each chronic condition corresponded to an average of approximately 0.24 hospitalizations per year. A similar pattern was observed for bed days (Fig. 1B). For people with five or fewer chronic conditions, the utilization of bed days was approximately proportional to the number of conditions. However, between five and six or more chronic conditions, a steep increase in utilization of bed days was observed. Among individuals with six or more chronic conditions, utilization was higher than the mean number of bed days per condition multiplied by the number of conditions.

**Fig 1.**
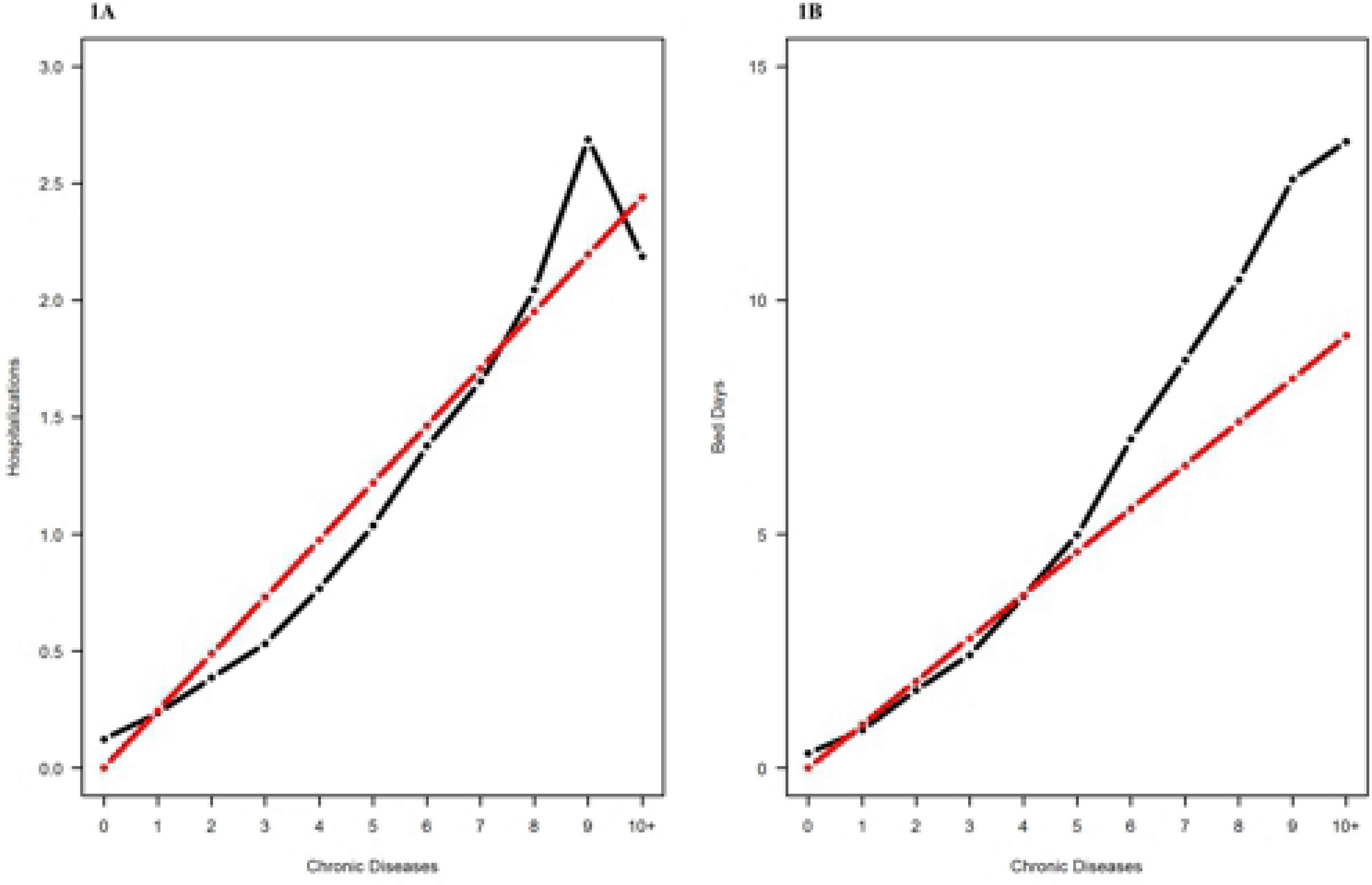
1A. Relationship between numbers of chronic conditions and hospitalizations. The black line indicates the number of hospitalizations by the number of chronic conditions. The red line indicates the mean number of hospitalizations multiplied by the number of chronic conditions. **1B. Relationship between numbers of chronic conditions and bed days.** The black line indicates the number of bed days by number of chronic conditions. The red line represents the mean number of bed days multiplied by the numbers of chronic conditions.

Fig 2 and Fig 3 depict utilization of hospitalization and bed days stratified by educational attainment. An educational gradient in hospitalization rates was observed across one to six or more chronic conditions (Fig 2). Hospitalizations were more frequent in individuals with shorter education, compared with those with longer education. For bed day utilization (Fig 3), individuals with no education exhibited the highest estimated utilization rates, regardless of the number of chronic conditions.

**Fig 2.**
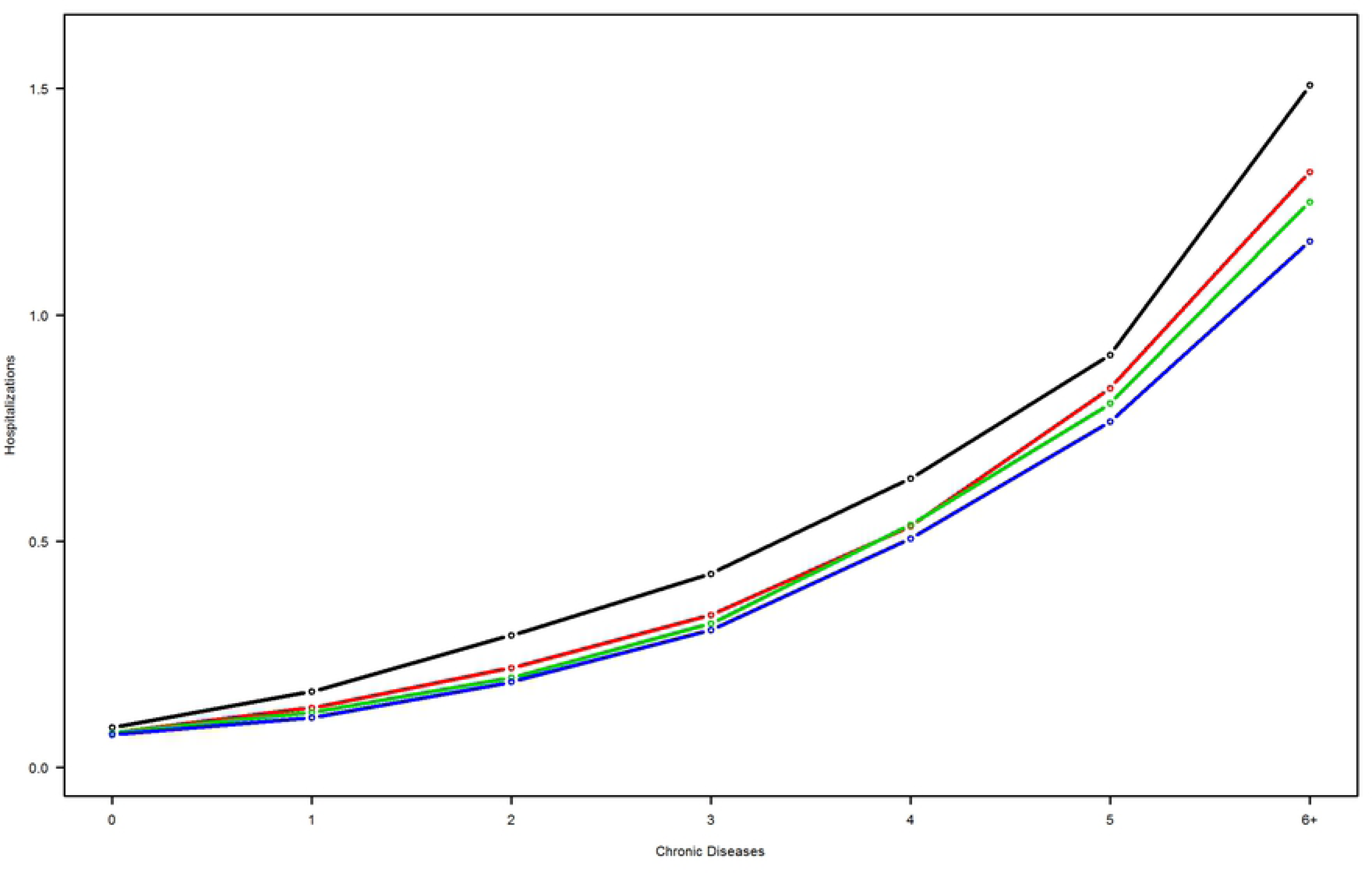
Number of chronic conditions associated with the modeled rate of hospitalizations by educational attainment levels. Black line, no education; red line, short education; green line, medium education; blue line, long education.

**Fig 3.**
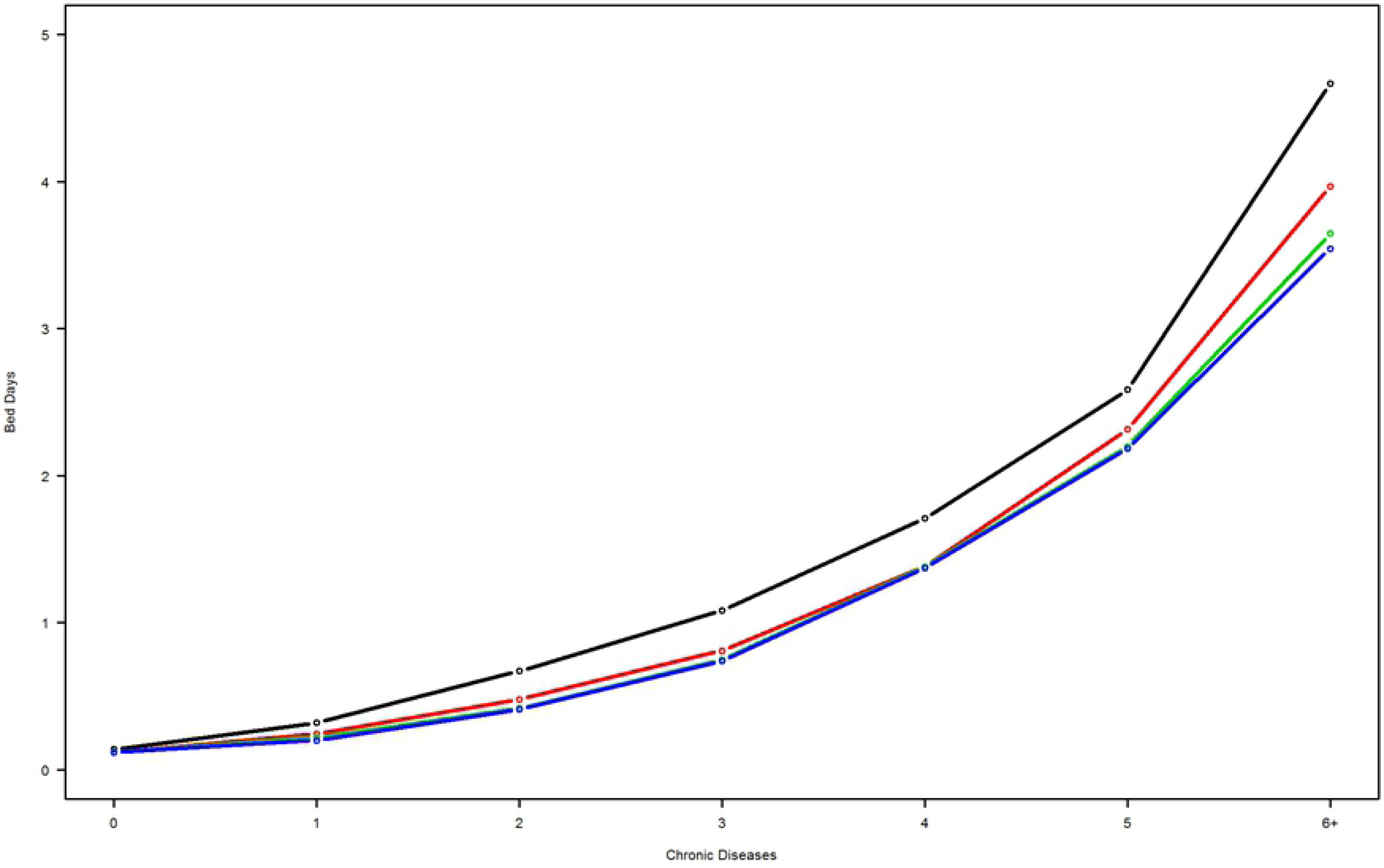
Number of chronic conditions associated with the modeled rate of bed days by educational attainment. Black line, no education; red line, short education; green line, medium education; blue line, long education.

## DISCUSSION

This study is the first large-scale, register-based study investigating associations between multimorbidity and utilization of healthcare services in the secondary and primary sectors in Denmark. In this study we were able to obtain a medical diagnosis for all diseased adults in the Capital Region aged 16 and above using algorithmic diagnoses for 16 conditions. This contrasts with studies using register-based diagnoses that only included patients who had had a hospital admission or were affiliated with an outpatient clinic. Although the 16 selected conditions do not represent the full spectrum of chronic disease, they include highly prevalent chronic conditions. A similar study from Scotland [33] included 40 conditions and revealed an age-stratified pattern of prevalence of chronic conditions nearly identical to that of the data reported here [7, 33]. This suggests that the 16 included diagnoses in this study encompass all conditions and combinations of conditions that are predominant at the population level.

The prevalence of multimorbidity in our study was 22% [7]. Females had a 10% overrepresentation among individuals with multimorbidity (Table 2). However, in this study the overrepresentation of women among multimorbid individuals cannot be explained by the longevity of women alone: The frequency of multimorbidity within each one-year age group was consistently higher for women than men from 68 years to 100 years of age, the difference increasing with age. This finding is consistent with previous results across healthcare systems and geography [34-36]. Eighty percent of the study population and 45% of individuals with multimorbidity were younger than age 65 (Table 2). The study of multimorbidity is often confined to adults aged 65 and up [19, 37-39]. However, younger individuals with multimorbidity who survive will age into this group and can be expected to have higher lifelong healthcare utilization. In fact, we observed that, compared to age-similar individuals without chronic conditions, younger multimorbid individuals had a relatively higher rate of bed day utilization than did older individuals with multimorbidity This finding highlights the importance of understanding chronic illness and healthcare utilization of younger multimorbid persons.

### Utilization vs. number of conditions

The consistency of increases in healthcare utilization for individuals with multimorbidity, compared to those without chronic conditions, is remarkable (Table 3). The use of all services was higher to a statistically significant degree. Excluding hospitalizations and bed days, the increase in utilization rates for individuals with multimorbidity ranged from 73% (emergency department visits) to 324% (outpatient visits), except for yearly control visits in general practice, which were 867% higher. Disregarding the latter due to very low utilization rates, the average increase in healthcare utilization was 180% for individuals with multimorbidity, compared to those with no chronic conditions. The corresponding average increase in utilization rates for multimorbid individuals, compared with those with one chronic disease, was 65%. Unadjusted ORs for healthcare utilization for individuals with multimorbidity, compared with those with no chronic conditions, were all statistically significant and ranged from 1.65 (95% confidence interval [CI], 1.63 – 1.66) for emergency department visits and 11.76 (95% CI, 11.58 – 11.96) for yearly control visits. When adjusted for gender, age, and educational attainment, utilization differences were less pronounced, ranging from 1.86 (95% CI, 1.83 – 1.88) for emergency department visits to 6.70 (95% CI, 6.64 – 6.76) for GP visits. A similar but less pronounced pattern was seen when comparing multimorbid individuals to those with one chronic disease (Table 3). These results demonstrate that the impact of multimorbidity on healthcare utilization applies to a range of services and varies relatively little. Healthcare utilization rates were 2-4 times higher than those for people without chronic conditions. The direct relationship between the number of chronic conditions and healthcare utilization is well-documented in the literature [14, 23, 24, 40, 41], but, to the best of our knowledge, the impact of multimorbidity on a broad spectrum of healthcare utilization has not been documented previously.

The approximately linear relationship between frequency of hospitalizations and the number of chronic conditions depicted in Fig. 1A also applies to the relationship between bed days and the number of chronic conditions in Fig. 1B for five or fewer conditions. When the number of conditions increases to six or more, the utilization of bed days increases by a factor greater than the impact of a single additional condition. The overall utilization pattern is that each condition corresponds to 0.24 hospitalizations and 0.92 bed days, while the length of each hospitalization is longer for individuals with six or more conditions. The slope of the curve for six or more conditions increases twofold to 1.84 (Fig 1B). The fact that the regression coefficient no longer explains the frequency of bed days per chronic condition leads us to define individuals with six or more conditions as high utilizers; they make up 0.87% of the population but account for 7.29% of bed days. An earlier Danish study found that 5% of the population with chronic conditions accounted for 45% of healthcare expenses [42]. This finding is in line with a large US study that showed that 5% of high utilizers accounted for up to 47% of healthcare costs [43]. A recent German study reported two subgroups of high utilizers: the oldest patients who suffered from severe multimorbidity and younger elderly patients with psychiatric or psychosomatic conditions [40]. Future research is required to examine characteristics and utilization of the high utilizers identified in this study.

Fig 1A shows a decline in utilization between nine and ten or more conditions. A possible explanation is that individuals with a very high disease burden have higher mortality, while those who survive have lower healthcare utilization than expected. Only 43 individuals had ten or more diagnosed conditions, and small sample size effects may also contribute to this finding.

### Socioeconomic status

We found that the prevalence of multimorbidity decreased with increasing educational attainment, revealing a pronounced and statistically significant inverse socioeconomic gradient. This is consistent with previous findings [7].

To study the impact of multimorbidity on healthcare costs, we adjusted data for proxy costs for different educational attainment groups with varying profiles in terms of age, gender, other healthcare utilization and level of multimorbidity. When adjusted for these effects, as described in the methods statistical section, a clear inverse social gradient in hospitalization utilization appeared (Fig. 2). Adjusted hospitalization utilization decreased as the level of educational attainment increased, generally irrespective of the number of chronic conditions. However, adjusted bed-day utilization revealed a slightly different pattern. An educational gradient no longer appeared; curves for short, medium, and high educational attainment tended to overlap. However, the group with no educational attainment stood out by virtue of higher healthcare utilization than the other. One important feature is that the relative difference between utilization rates for individuals with no education and those in other educational groups appeared close to constant across the number of chronic conditions.

In general, the number of hospitalizations decreased with increasing level of educational attainment. For multimorbid individuals, the length of each hospitalization was longer for individuals without any education than for those with at least some education. The reason for this finding is unknown. It may be the case that chronic disease tends to be more severe among individuals without education and that longer hospitalizations are caused by non-chronic conditions not included in our study. In addition, one could posit that different spectra of chronic conditions occur for persons with and without education, but investigation of this supposition will require further research. Other factors affecting the higher healthcare utilization rate observed in people with lower education attainment include lower health literacy levels linked to lower education attainment [44]. Disease burden has been shown to be associated with lower education levels in type 2 diabetes, and this may apply to other chronic conditions [45]. Furthermore, people with none or low educational attainment often has weak social networks and proper discharge to home might demand coordinated preparation that rely, in part, on support from individual’s own social support structure [46]. However, we currently lack a well-founded explanation for differences in the two adjusted proxies for general utilization of healthcare services.

### Strengths and limitations

The major strength of our study is that it is a large-scale register-based study, including comprehensive information about chronic conditions, healthcare utilization, and educational attainment of the complete population of the Capital Region of Denmark aged 16 years and above. Generally, data from the Danish national registers provide complete information about healthcare system contacts, are of high quality and reliability, and are used extensively in research [25, 47]. As a general population-based study, our findings reflect the actual situation in a real-world setting. Based on register data, our study was free of recall bias and there was no loss of follow up.

Several limitations of our study deserve consideration. The necessary use of diagnostic algorithms to identify patients with chronic conditions in the primary healthcare sector is an approximation of actual diagnoses. Although the diagnostic algorithms have been shown to be highly accurate [6], they are not clinically determined by physicians. The study was based on cross-sectional data collected during a single year, and the number of patients with specific chronic conditions may be underestimated [48, 49]. In addition, we were not able to include healthcare services provided in the municipalities. Scarce register data exist for municipality services; existing data are not well defined and thus not useful for research purposes. Inclusion of the data on utilization of community healthcare services would likely have helped to generate a more complete picture of healthcare utilization related to multimorbidity. Finally, 6% of our population did not have information on educational attainment. This information appeared to be missing at random, except for individuals aged 91 years and above; Danish administrative registers only contain information on education for individuals born since 1921 [31]. However, this group contained few individuals, and we estimated the effect of these missing data to be minimal and possible changes in parameter estimates therefore very small.

Comparison to other studies on multimorbidity in populations should be performed with care. Varying definitions of multimorbidity (i.e., two or more chronic conditions), included conditions, and data collection methods render comparisons difficult. However, these challenges may be overcome for large studies [33].

## Conclusion

Multimorbidity is associated with a significant increase in utilization of all healthcare services in Denmark. A socioeconomic gradient was observed in utilization of hospitalizations, and socioeconomic effects in utilization of bed days. A steep increase in the utilization of bed days in patients with six or more chronic conditions suggests a subpopulation of high utilizers that should be explored in further studies.

## Acknowledgements

We thank Jennifer Green for skillful editing.

